# Development of efficient RNAi methods in the corn leafhopper *Dalbulus maidis*, a promising application for pest control

**DOI:** 10.1101/2022.01.17.476645

**Authors:** L.I. Dalaisón-Fuentes, A. Pascual, E. Gazza, E. Welchen, R. Rivera-Pomar, M.I. Catalano

## Abstract

**BACKGROUND:** The corn leafhopper *Dalbulus maidis* is the main vector of three important stunting pathogens that affect maize production. The most common control strategy against this species is the use of insecticides that provide minimal, short-term protection. In this context, genomic-based technologies such as RNA interference (RNAi) could be a suitable approach to control this pest in a highly specific manner, avoiding the adverse effects associated with insecticide misuse. Therefore, the objective of the present work was to assess the application of RNAi on *D. maidis* through different dsRNA delivery methods and known the function of target gene, *Bicaudal C* (*BicC*).

**RESULTS:** We have identified and characterized the core components of the RNAi machinery *in silico* and established two methods of exogenous double-stranded RNAs (dsRNA) delivery to *D. maidis*. *BicC* -an important regulator of insect oogenesis-dsRNA was successfully delivered via injection or ingestion to adult females, causing significant reductions in the transcript levels and ovipositions and observable phenotypes in the ovaries when compared to control females. The small doses of *dsRNA^BicC^* administered were enough to trigger a strong RNAi response, demonstrating that *D. maidis* is highly sensitive to RNAi.

**CONCLUSION:** This is, to our knowledge, the first report describing RNAi application in *D. maidis*, a tool that can be used to advance towards a novel, insecticide-free control strategy against this pest.

## 1. INTRODUCTION

The corn leafhopper *Dalbulus maidis* (DeLong & Wolcott) (Hemiptera: Auchenorrhyncha) is the main vector of maize rayado fino virus (MRFV)^1^, maize bushy stunt phytoplasma (MBSP) and *Spiroplasma kunkelii*^2^ that affect maize (*Zea mays* L.) crops throughout the Americas^3^. *D. maidis* transmits these pathogens, either alone or in combination, in a persistent-propagative mode^4,5^, causing the diseases referred to as Corn Stunt^2,4,5^. High infestation rates, which can reach 100% in some areas, may result in yield losses of up to 90%, representing a relevant threat to agriculture^6,7^.

Current control strategies against *D. maidis* are limited to corn resistant germplasm^8^ and use of insecticides as seed treatments or foliar sprays^9–11^. While insecticides applied to seed treatment only protect the plant at early growth stages, foliar insecticides usually fail to reduce maize damage and incidence of stunting diseases^12,13^, additionally, they cannot discriminate between pest and non-pest species, and widespread use can lead to resistance to them^14–16^. In this context, RNA interference (RNAi) emerges as a promising alternative for vector control^17^.

RNAi is a highly conserved, sequence-specific mechanism^18^ that has been widely used for elucidating gene function, especially in non-model organisms^19^ and in recent years, it has become a powerful tool for insect pest management^20–22^. Briefly, RNAi mechanism consists of two main events: first, exogenously delivered double-stranded RNAs (dsRNA) are internalized and processed by the cellular RNAi machinery into small interfering RNA (siRNA), which in second turn triggers the degradation of complementary endogenous messenger RNA (mRNA) leading to gene silencing^23–25^. The core components comprising the RNAi machinery have been identified and characterized in several hemiptera species^26–33^, but the mechanism underlying dsRNA internalization and signal spreading (i.e., systemic RNAi) still remains uncertain^34^.

Most RNAi research has been carried out via microinjection and natural/artificial diets ^35–37^, and by detached leaf/petiole dip, topical application to either the insect or plant or transgenic plants expressing insect-derived dsRNAs^20–22^, to a lesser extent. Besides the route of entry, RNAi effectiveness has been found to be highly variable among insect species^38^. These variations mainly depend on intrinsic factors of the species, being some orders more refractory to dsRNA administration than others^39–41^, as well as extrinsic factors, including the delivery method and the target gene^42^. However, gene silencing has been successfully achieved in several hemipterans, including the leafhoppers *Amrasca biguttula*^43^, *Circulifer haematoceps*^44^, *Graminella nigrifrons*^45^, *Homalodisca vitripennis*^46,47^ and *Nephotettix cincticeps*^48–50^, along with others.

Due to the central role of reproduction on insect life cycle and propagation, we decided to target *Bicaudal C (BicC)*. It has been demonstrated that *BicC* plays a vital role during oogenesis in other hemipterans^51,52^ and in the dipteran *Drosophila melanogaster*, in which is also responsible for specifying anterior-posterior (AP) polarity of the embryos^53–55^. Here we silenced this gene in order to assess the application of RNAi in *D. maidis.* We have identified and characterized core components of the RNAi pathway and established two reliable methods of dsRNA delivery. We also describe the essential function of *Dmai-BicC* in female reproduction, being the first time that this species has been studied from a genetic point of view. Our findings enable further molecular analyses on this leafhopper and provide a framework for developing new control strategies.

## 2. MATERIALS AND METHODS

### 2.1. Insect rearing

A healthy colony of *D. maidis* was maintained on corn plants (*Zea mays* L.) in our laboratory. The colony was kept in aluminum-framed cages with a fine *voile*-type nylon mesh and placed in a greenhouse at a temperature of 25°C and 80% relative humidity, with a photoperiod of 16:8 hours (h) (light: darkness). At 23±3°C, embryogenesis is completed 11.5±1.3 days after egg laying^56^.

### 2.2. Evaluation of feeding behavior on artificial diet

To test whether the insects would feed on the artificial diet containing the dsRNAs, a liquid artificial diet consisting of 10% (w/v) sucrose solution^57^ was prepared by dissolving sucrose in sterile water on a magnetic stirrer hotplate (25°C) and then, green food coloring (Fleibor S.R.L) was added. This mixture was held between two layers of stretched Parafilm M (Bemis™) located at the top end of the feeding chamber. The other end was covered with a fine nylon mesh to allow aeration. Insects were placed inside the feeding chamber during 3-5 h. Chambers were checked during the following days. Electrical penetration graph (EPG) technique was used to corroborate feeding on the artificial diet^58,59^. The feeding platform was constructed from a small plastic Petri dish according to Trębicki *et al.*^60^. Insects were monitored for 8 h in a Giga-8 EPG model (EPG Systems, Wageningen, The Netherlands).

### 2.3. Identification of RNAi-related genes and *Dmai-BicC*

Gene identification was performed using local BLAST^61^ on a *D. maidis* adult transcriptome, previously assembled in our lab^62^ (**File S1**). The search of the RNAi machinery genes was limited to *Dicer2 (Dcr2), Argonaute2 (Ago2), R2D2 (R2D2)* and *Systemic RNA interference defective protein 1 (Sid1)* proteins from other hemipterans available in the NCBI protein database. Best hits were chosen considering the percent identity (%ID), match length and E-value (≤1E-5). ORF finder tool^63^ was used to detect open reading frames (ORFs) in transcript sequences. Protein signatures were predicted by InterProScan (version 5.52-86.0)^64^. Translated sequences were used as a query to perform BLASTp^65^ searches against the non-redundant protein database (version 1.1, accessed September 2021). To provide additional confirmation on identity, each sequence was aligned with orthologues from other species using Clustal Ω (version 1.2.4)^66^. Conserved transmembrane helices found in *Sid1* proteins were predicted for *Dmai-Sid1* at TMHMM Server (version 2.0)^67^. The same workflow was applied for *BicC*.

### 2.4. dsRNA synthesis

Total RNA was isolated from ovaries from adult females using TRIzol™ reagent (Invitrogen), according to manufacturer’s instructions. cDNA was synthesized following the EasyScript Reverse Transcriptase (AP-Biotech) protocol and used as a template for RT-PCR. Specific primers for *Dmai-BicC* were designed^68,69^ to amplify a region of 372 bp **(Table S1)** within one of the predicted KH domains **(Fig. S1)**. Reaction conditions were 2 minutes (min) at 94°C, followed by 35 cycles of: 92°C for 30 seconds (s), 56°C for 30 s and 72°C for 45 s, and a final extension step for 4 min at 72°C (Taq Pegasus, Productos Bio-Lógicos). The integrity of the amplicon was checked in a 1% agarose gel and sequenced to confirm its identity (Macrogen Inc.).

The dsRNA targeting *Dmai-BicC* (*dsRNA^BicC^*) was synthesized using T7 Polymerase (ThermoFisher), according to the manufacturer’s specifications. The same sense and antisense primers were designed containing T7 promoter sequence at the 5’end for further use during *in vitro* transcription **(Table S1)**. dsRNA was purified by DNase digestion (Qiagen). Purified dsRNA was examined in a 1% agarose gel to ensure its integrity. A negative control was included using the *β-lactamase* gene (*dsRNA^βlac^*) amplified from a pRSET b plasmid^70^. The final doses of dsRNA were 100 ng/μL (*dsRNA^BicC100^*) and 200 ng/μL (*dsRNA^Bic200^*) for *dsRNA^BicC^* and 130-170 ng/μL for *dsRNA^βlac^* treatments.

### 2.5. dsRNA feeding

Newly molted adult females were anesthetized with CO_2_ and placed inside the feeding chambers where they fed only artificial diet. 24 h later, the dsRNA was added as a supplement to the diet. Insects fed by puncturing the inner membrane of the diet pouch. This solution was renewed daily for three consecutive days. After three days of dsRNA ingestion, each female was placed individually in a cage and mated with one non-exposed male, allowing them to feed and lay eggs on corn plants. Female mortality and oviposition (i.e., number of eggs laid per female) were recorded every 48 h. Rearing conditions were the same as for the colony.

### 2.6. dsRNA microinjection

Microinjection was performed with a glass needle (P-30 model, Sutter Instrument Company) between two abdominal sternites. Micropipette puller settings were the following: 950 (heat #1), 950 (pull) and 3.24-3.25 (optical micrometer). Newly molted adult females were anesthetized with CO_2_ and placed on a frozen surface, to ensure their lethargy throughout the procedure. After microinjection, the females were placed in individual cages attached to maize plants to allow feeding. 24 h later, once the females were recovered from injury, one male was introduced into each cage. From this stage on, the experiment continued as described for dsRNA feeding.

### 2.7. Ovary and embryo manipulation

After 18 days since dsRNA administration, ovaries were dissected according to Pascual *et al.*^52^, but fixation step was limited to 10-15 min.

Ovipositions were monitored during the expected time of embryogenesis^56^ and eggs which did not hatch were fixed as follows: they were collected using insect pins under a stereomicroscope (Zeiss, Stem 305), placed in microtubes and incubated at −80°C for 3-5 min. Then 200 μL of PBS 1X were added and submerged into hot water (85-90°C) for 1 min. Afterwards, 60 μL of 37% Formaldehyde (FA) were added to the PBS 1X and fixed for 10-15 min on a shaking platform (150-200 rpm). Subsequently, 260 μL of heptane were used to post-fix the eggs for 10-15 min on a shaking platform at the same speed. After the two phases separated, the upper phase was removed and the lower phase was rinsed three times with 500 μL of 100% Methanol (MeOH) followed by high speed vortex. Finally, the chorion and perivitelline membrane of the eggs were manually removed under a stereomicroscope (Zeiss, Stem 305).

For fluorescence microscopy, fixed ovaries and eggs were incubated in a 300 nM DAPI stain solution (4′,6-diamidino-2-phenylindole, ThermoFisher Scientific) on a shaking platform (150-200 rpm) for 15-20 min. Prior to staining, eggs were rehydrated through the subsequent series of MeOH in PBS: 75% MeOH / 25% PBS 1X, 50% MeOH / 50% PBS 1X and 25% MeOH / 75% PBS 1X, for 3-5 min in each solution. Images were acquired with a stereomicroscope and a fluorescence microscope (Zeiss, Axio Imager A2).

### 2.8. Real-time quantitative PCR (RT-qPCR)

Independent experiments were conducted to determine *Dmai-BicC* mRNA expression after dsRNA delivery. We decided to evaluate the highest dsRNA concentration (200 ng/μL). Total RNA samples were prepared from ovaries dissected at 1- and 7-days, based on oocyte maturation^71,72^, after completion of dsRNA administration. Ovaries were dissected in PBS 1X and immediately stored in TRIzol™ reagent (Invitrogen) for RNA extraction. RNA was quantified by UV-Vis spectrophotometric measurement in a Nanodrop 2000c Instrument (Thermo Scientific™). Concentration and purity ratios (A260/280 and A260/230) were analyzed. Total RNA of individual replicates was subjected to DNase I, RNase-free (Thermo Scientific™) treatment according to the manufacturer’s indications. First strand cDNA synthesis was performed using the logo (dT)_18_ primer and RevertAid Reverse Transcriptase (Thermo Scientific™) according to the protocol provided by the manufacturer. Real time (RT)-qPCR was carried out in a Bio-Rad® CFX-96™ thermocycler in 10 μl final volume reaction using SsoAdvanced™ Universal SYBR® Green Supermix (Bio-Rad) and 50 pmol of each forward and reverse primer **(Table S1)**. Thermal cycle protocol consisted of 30s at 95°C for initial denaturation, followed by 35 cycles of 15s at 95°C for denaturation and 20s at 60°C for annealing and extension. Melting curve analysis was performed at the end by using instrument default settings (65°C-95°C, 0.5°C temperature increment). Relative transcript levels were calculated by a comparative Ct method^73^. Expression values were normalized using *Actin 1* (*Dmai-Act1*, **Table S1**), after a screen of several housekeeping-gene candidates, as it provided consistent results on the samples analyzed (data not shown).

### 2.9. Statical analysis

All statistical analyses were performed using the InfoStat^74^ statistical software. A generalized linear mixed model (GLMM) was applied for the number of eggs laid to model negative binomial variables, being “treatment” the fixed effect and “female” the random effect. The model was adjusted using nlme^75^ and lme4^76^ packages from R language^77^. Predicted values were compared using the DGC test^78^ with a significance level of 5%. Proportions of surviving insects were calculated using the Kaplan-Meier method and the p-values generated via Log-rank test were used to test the null hypothesis that the survival curves were identical in the three treatment groups, with a significance level of 5%.

## 3. RESULTS

### 3.1. *D. maidis* feeding on artificial diet

We confirmed that *D. maidis* adults can survive *in vitro* for at least five days by feeding exclusively on artificial diet **(Fig. 1A-B)**. Feeding was evidenced by green honeydew droplets **(Fig. 1C)**, as a result of the green food coloring included in the diet. Furthermore, insects presented the same green coloration in their mouthparts **(Fig. 1D)** and abdomen **(Fig. 1E)**, proving that the diet had been ingested and passed through the gut. Furthermore, EPG recordings obtained from *D. maidis* having access to the artificial diet produced the typical waveform correlated with active feeding **(Fig. 1F)**. These results indicate that *D. maidis* feeds reliably on the artificial diet and, therefore, can acquire the dsRNA orally.

**Figure 1.**
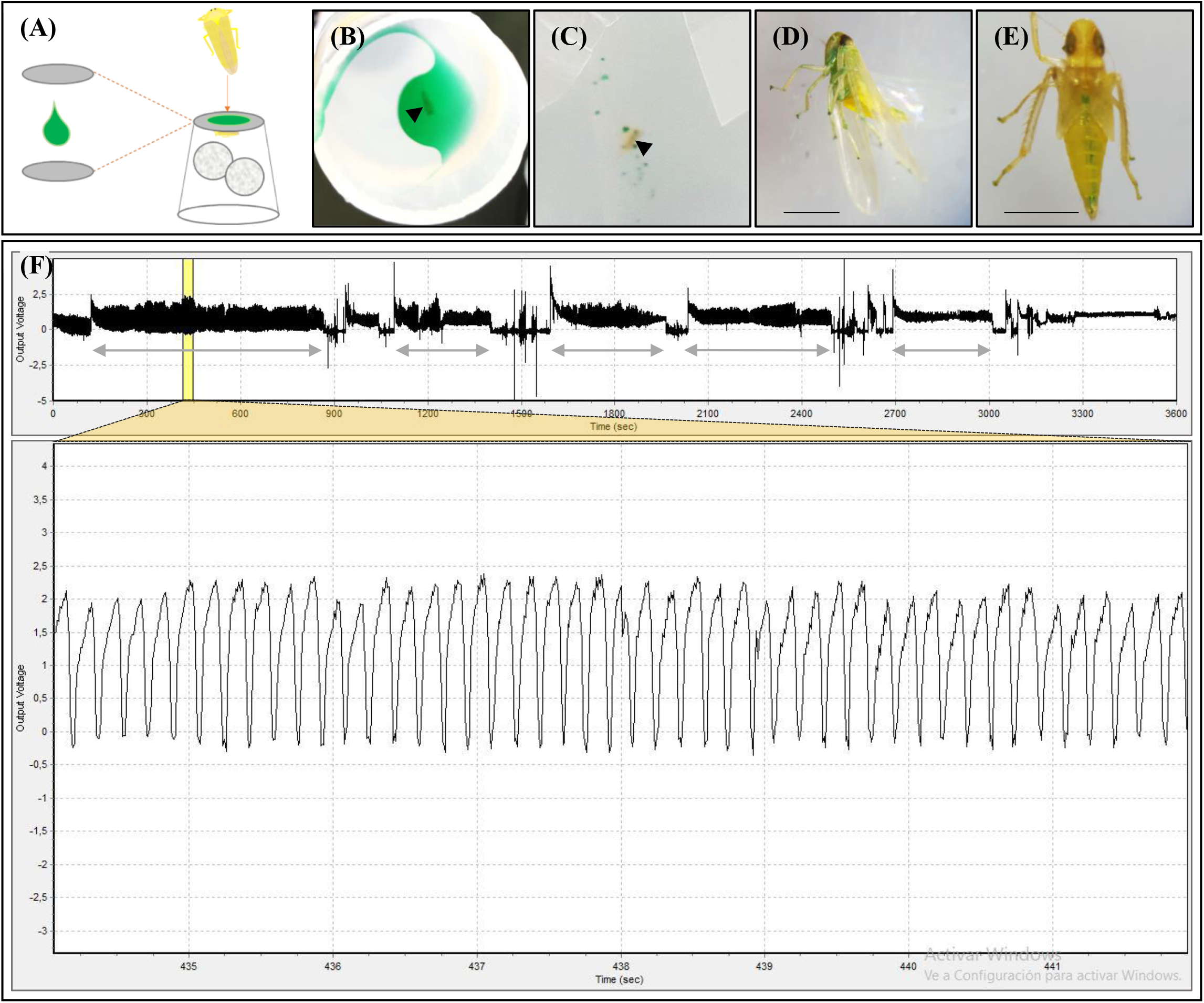
Feeding behavior of *D. maidis* on the artificial diet. **(A)** Schematic representation of the green food coloring assay. **(B)** Colored diet located at the top end of the feeding chamber. **(C)** Green honeydew droplets on the wall of the feeding chamber. **(D, E)** Green-colored mouthparts and abdomen, respectively. Scale bar: 1 mm. The black arrowheads in **(B)** and **(C)** point to *D. maidis* adults. **(F)** Upper graph: EPG waveform of *D. maidis* having access to the artificial. Double-headed arrows indicate longest time periods of active feeding. Lower graph: 8 s of active feeding corresponding to the yellow highlighted region of the upper graph. X axis: time (s), Y axis: voltage.

### 3.2. *In silico* identification of core RNAi machinery genes on *D. maidis*

We identified the active expression of the core components of RNAi machinery on *D. maidis* adult transcriptome (**Table 1)**. *Dmai-Dcr2*, *Dmai-Ago2*, *Dmai-R2D2* and *Dmai-Sid1* translated sequences all showed the protein signatures characteristic of the corresponding families previously described **(Fig. S2)**. Sequence similarity searches against the protein database revealed high similarity with orthologs from closely related insect species, providing further confirmation of their identity (data not shown). Altogether, these results demonstrate that the genes involved in RNAi (siRNA) pathway are conserved in *D. maidis*.

**Table 1.** Identification of the genes involved in the activation of the silencing machinery and dsRNA uptake in *D. maidis.*

| Annotation | Transcript ID | cDNA(bp) | ORF | EPL (aa) | BLASTx |  |  |  |
| --- | --- | --- | --- | --- | --- | --- | --- | --- |
|  |  |  |  |  | Best hit | % ID | Match length | E-value |
| ***Dcr-2*** | TRINITY_DN24319_c0_g1_i1 | 5.719 | 81-5.084 | 1.667 | *Nc* (APF46959.1) | 63,19 | 1.668 | 0.0 |
| ***Ago-2*** | TRINITY_DN25305_c1_g2_i2 | 5.267 | 5.168-2.451 | 905 | *Rd* (AMN92166.1) | 99,77 | 854 | 0.0 |
| ***R2D2*** | TRINITY_DN22775_c0_g2_i1 | 2.052 | 1.738-518 | 406 | *Nv* (AVK59464.1) | 37,77 | 188 | 1.97e-29 |
| ***Sid-1*** | TRINITY_DN25274_c0_g1_i2 | 4.365 | 3.954-1.603 | 783 | *Nl* (AGH30336.1) | 44,51 | 737 | 0.0 |

Annotations were assessed by sequence similarity search. cDNA: transcript length; ORF: Open Reading Frame; EPL: Encoded Protein Length; Best hit: the closest match found by local BLASTx searches; % ID (percent identity), match length and e-value to the best hit. *Nc*: *Nephotettix cincticeps*, *Rd*: *Recilia dorsalis*, *Nv*: *Nezara viridula* and *Nl*: *Nilaparvata lugens*.

### 3.3. Effect of RNAi on female survival and fertility

To determine the efficiency of RNAi gene silencing on *D. maidis*, female mortality and oviposition were recorded at 2-day intervals. No significant differences were found in the proportion of living females throughout the experiments **(Fig. S3)**, regardless of the delivery method employed (microinjection: p=0.9475 and feeding: p=0.1406).

For the microinjection assay, both doses of *dsRNA^BicC^* (100 and 200 ng/μL, *dsRNA^BicC100^* and *dsRNA^BicC200^*, respectively) reduced significantly the ovipositions when compared to the *dsRNA^βlac^* females, remaining close to zero throughout the experiment. Also, significant differences between both *ds*RNA^*BicC*^ concentrations were observed **(Fig. 2A)**. Similarly, both *dsRNA^BicC100^* and *dsRNA^BicC200^*-fed females had significantly lower ovipositions than those in the control ones; however, they laid more eggs than the *dsRNA^BicC^*-injected females. No significant differences between the *dsRNA^BicC^* concentrations were detected for the feeding assay **(Fig. 2B)**.

**Figure 2.**
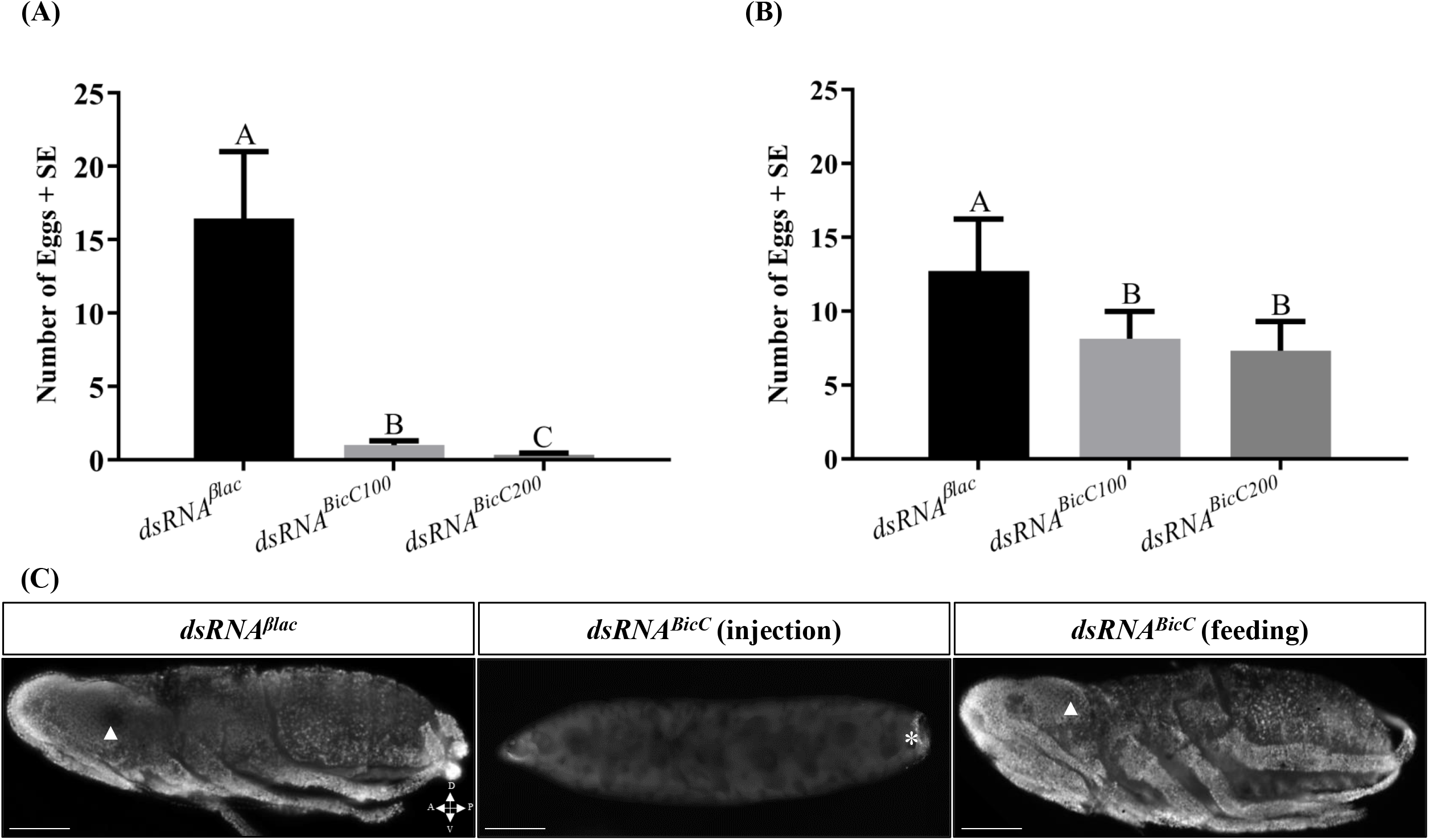
Ovipositions of *D. maidis* females after dsRNA delivery. Ovipositions in the injection- **(A)** and feeding **(B)**-based experiments. The number of eggs laid by the females showed a significant effect of the treatment factor (microinjection: p≤0.0001 and feeding: p=0.0026). Each bar represents the mean number of eggs laid by *D. maidis* females and the standard error of the mean (SE). Values with the same letter are not significantly different according to contrasts in the mixed model test (p≤0.05). **(C)** Nuclei distribution of the eggs laid by control and *dsRNA^BicC^*-treated females by DAPI staining. Scale bar: 100 μm. White asterisk: *D. maidis* endosymbiont. White arrowhead: embryos’ eye. A: anterior; P: posterior; D: dorsal; V: ventral.

The few eggs laid by the *dsRNA^BicC^*-injected females compared to those in the control **(Fig. 2A and S4A)** were smaller, thinner and had abnormally large bubbles in the yolk **(Fig. S4B)**. None of them hatched and had no distinguishable embryonic structures, indicating that embryogenesis was affected **(Fig. 2B)**. Unlike *dsRNA^BicC^*-injected females, most of the eggs laid by *dsRNA^BicC^*-fed females reached first-instar larvae within the expected time **(Fig. 2C and S4C)**. These results indicate that microinjection and ingestion of dsRNA can successfully trigger an RNAi response in *D. maidis*.

### 3.4. Effect of RNAi on *Dmai-Bic* transcript levels

Delivery of *dsRNA^BicC^* by microinjection resulted in significant reductions in *Dmai-BicC* transcripts when compared to the *dsRNA^βlac^* control group (p≤0,005). The strongest silencing of *Dmai-BicC* was measured at day 7 after dsRNA administration, with a decreasing of a fold change near to 3.2 times at 1 day after treatment and more than 170 times at day 7 in *dsRNA^BicC^*-injected females in comparison to the expression in control ones **(Fig. 3A)**. Significant differences in the abundance of *Dmai-BicC* transcripts were also detected between the females fed with *dsRNA^BicC^* and those with *dsRNA^βlac^* (p≤0,005) at 1 day and 7 days after dsRNA delivery, decreasing 2.6 and 3.7 times respectively **(Fig. 3B)**. A greater and faster reduction in *Dmai-BicC* transcript abundance was obtained by microinjection rather than feeding.

**Figure 3.**
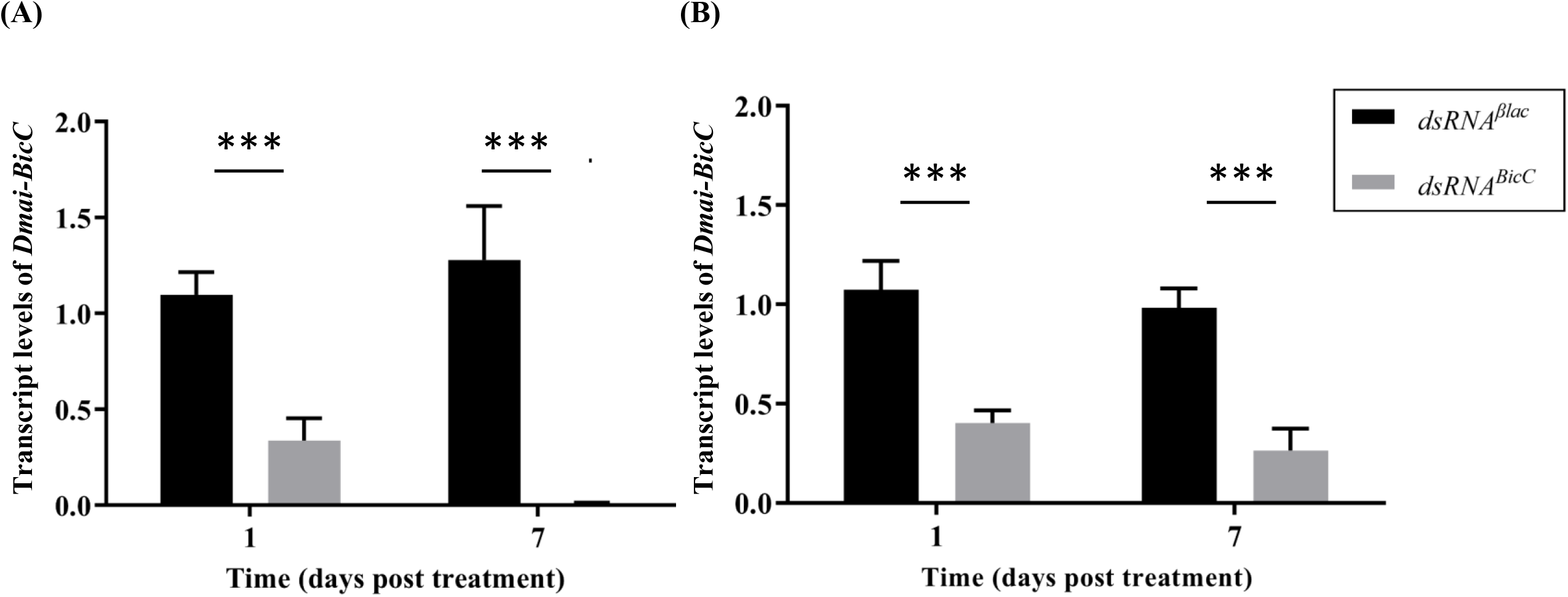
Expression levels of *Dmai-BicC* after dsRNA delivery. Transcript levels of *Dmai-BicC* during the progression of injection **(A)** or feeding **(B)**-based experiments. Expression was measured at 1 and 7 days after administration of *dsRNA^BicC^* (gray bars) or *dsRNA^βlac^* (black bars) used as an experimental control. All values were referred to as the *Dmai-BicC* expression at 1-day post-treatment in females treated with *dsRNA^βlac^*. Each bar represents the mean and standard deviation (SD) of three biological replicates. *** p≤0.005 (ANOVA, LSD Fisher test).

### 3.5. *Dmai-BicC* role during oogenesis

Morphological examination of the ovaries under stereomicroscope showed that 87,5% of the females injected with *dsRNA^BicC100^* presented the phenotype in **Fig. 4B**. The rest of them exhibited the same characteristics as *dsRNA^βlac^*-injected females **(Fig. 4A)**. This percentage increased to 100% for the group injected with *dsRNA^BicC200^*. Silenced females had the same number of tubular-shaped ovarioles -germarium and developing oocytes surrounded by follicle cells- as the controls, but retained a greater number of oocytes, as they failed to advance towards choriogenesis. In addition to their smaller size, the cytoplasm of the oocytes displayed a white coloration, in opposition to the translucent cytoplasm of control vitellogenic oocytes. Conversely, alterations in the ovaries of *dsRNA^BicC^*-fed females ranged from control (average of 79,7%) **(Fig. 4C and S5A)** to a variety of morphological defects (13.3% and 27.3% of the females fed with *dsRNA^BicC100^* and *dsRNA^BicC200^*, respectively) **(Fig. S5B-C)**.

**Figure 4.**
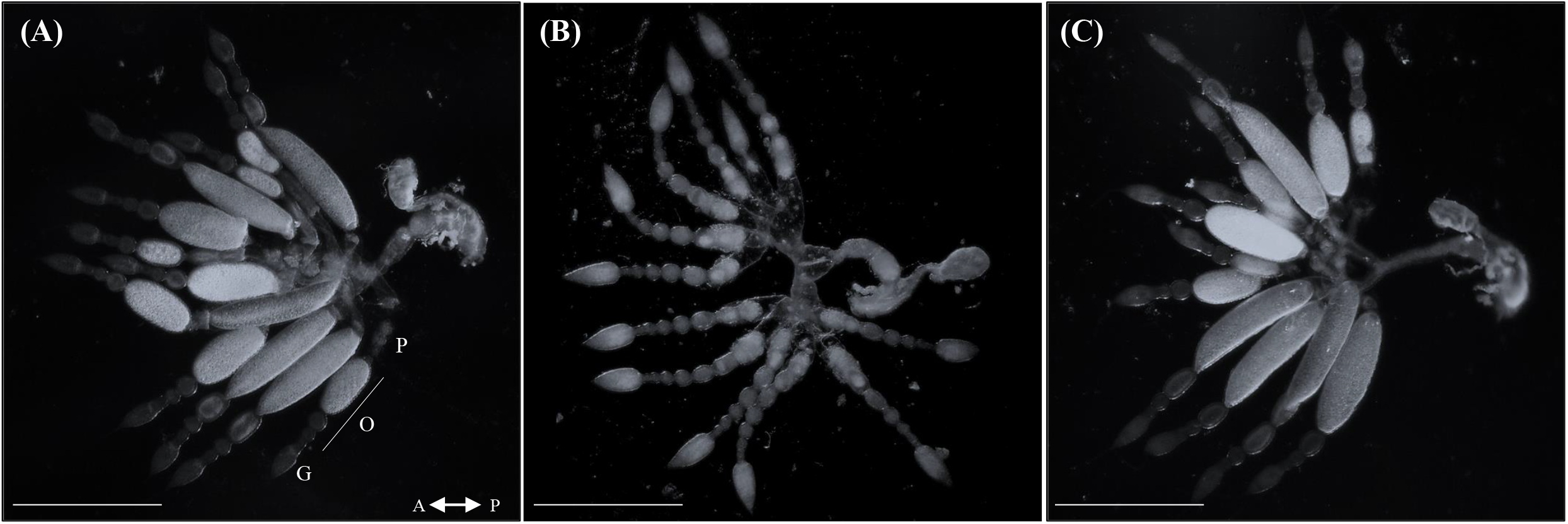
Effect of dsRNA on *D. maidis* female reproductive organs. Ovary morphology of a control **(A)** *dsRNA^BicC^*-injected **(B)** and *dsRNA^BicC^*-fed **(C)** female under dissecting microscope. Scale bar: 500 μm. G: germarium; O: oocytes and P: pedicle. A: anterior; P: posterior.

The ovaries dissected from injected females were analyzed in more detail by DIC optics and nuclei staining to determine cell distribution. We corroborated that *dsRNA^BicC^* did not produce changes in the germarium (data not shown). The follicular epithelium was present in the ovaries of silenced females, but as the oocytes advanced through the ovariole they got progressively more folded and wrinkled **(Fig. 5D)**, causing the follicular cells to lose the regular columnar shape seen in control females **(Fig. 5A)**. Likewise, nuclei became closer to each other to the point of overlapping, with no intercellular space left among them **(Fig. 5E-F)**.

**Figure 5.**
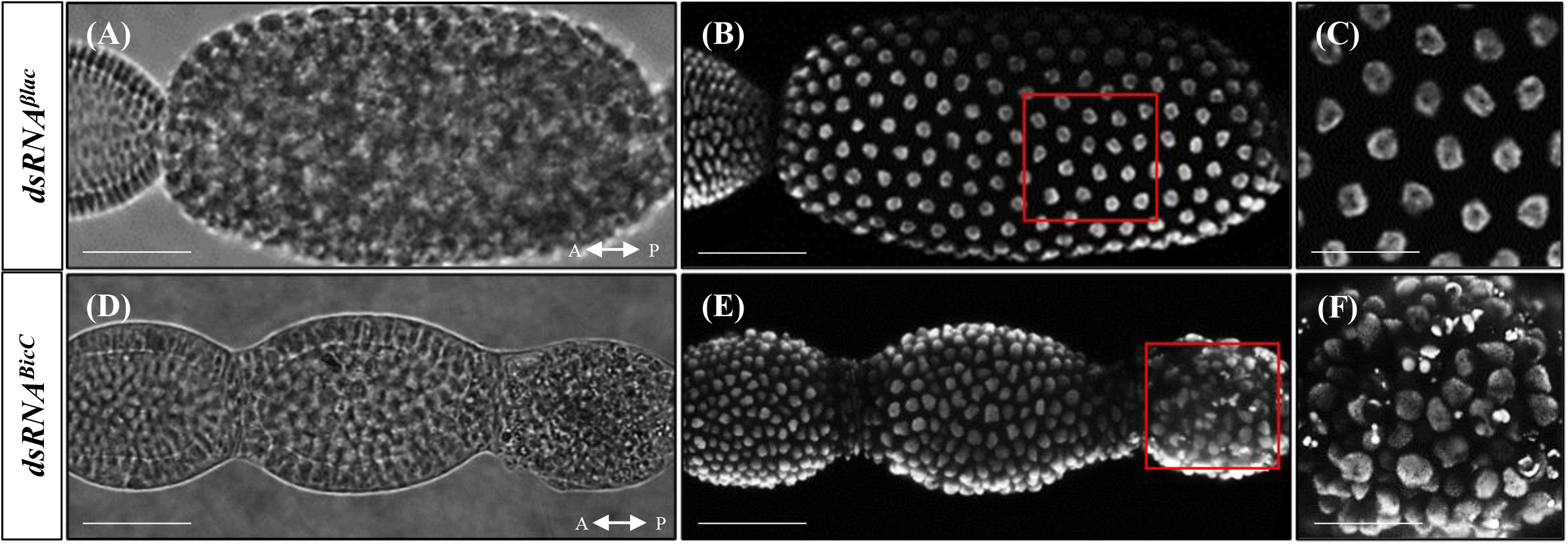
Effect of dsRNA on *D. maidis* follicular epithelium. Oocytes of control and *dsRNA^BicC^*-injected females by differential interference contrast (DIC) microscopy **(A, D)** and their nuclei distribution by DAPI staining **(B, E)**. Scale bar: 100 μm. **(C, F)** Follicular epithelium of control and *dsRNA^BicC^*-injected females showing the nuclei distribution of the highlighted regions in **(B)** and **(E)**. Scale bar: 50 μm.

These phenotypes agree with the observed reduction in the transcript levels and ovipositions and support the idea that the lack of *Dmai-BicC* affects oogenesis and oocyte maturation.

## 4. DISCUSSION

Developing new, RNAi-based control strategies is crucial in order to achieve specific pest control^79^. Finding suitable dsRNA delivery methods constitute the first step for the development of these control techniques^35,80^. This study presents, to the best of our knowledge, the first procedure for dsRNA delivery in *D. maidis* via microinjection and feeding and successfully silenced *Dmai-BicC*, an essential gene of oogenesis, thus providing with a potential tool to control this pest in a specific, sustainable way.

Core RNAi machinery genes -*Dcr2*, *Ago2* and *R2D2*-were found in *D. maidis* adult transcriptome, as have been reported in several hemipterans^26,27,29,30,32,33^, with the exception of *Halyomorpha halys*^28^ and *Diaphorina citri*^31^, in which *R2D2* was shown to be absent. We were also able to identify *Sid1*, a key component in the uptake and systemic spread of dsRNA^81^. *Sid-* like channel proteins have been characterized in many phytophagous species^26,27,29–31^, excluding some bug species^28,33^. The presence of these core genes in our transcriptomic database clearly indicates that the RNAi pathway is conserved in *D. maidis*.

In this context, we established an *in vitro* rearing system that allowed us to easily deliver the dsRNA. The addition of green food coloring served as proof of diet ingestion, using the same approach as Ghosh *et al.*^80^ for evaluating the potential of green beans to deliver dsRNA in the brown marmorated stink bug, *H. halys*. Parallel EPG recordings confirmed our observations and were consistent with previous studies in other leafhoppers, such as *Empoasca vitis*^82^ and *Orosius orientalis*^60^, where active feeding in artificial diets was demonstrated.

To assess the application of RNAi on *D. maidis*, we first attempted to silence a gene in adult insects that led to a visible non-lethal phenotype, but provides the opportunity to control this species by reducing the rate of population increase. Both methods -microinjection and feeding-significantly lowered transcript abundance and egg production, but differed in the morphological alterations generated in the ovaries. Our findings also revealed that *D. maidis* is sensitive to RNAi, as small doses of dsRNA (100-200 ng/μL) could induce strong RNAi responses, analogous to that reported by Taning *et al.*^31^ in the Asian citrus psyllid, *D. citri*.

The efficiency of oral delivery in knocking down the target gene was not as high as that accomplished by microinjection, in agreement with earlier research in the corn planthopper *Peregrinus maidis*^72^. These differences between delivery methods may be attributed to the lower amount of dsRNA delivered to *D. maidis* females by ingestion and/or the rapid degradation of dsRNA in the diet prior to ingestion, or in the digestive tract of the insects after feeding. In accordance with this, the presence of adverse pH conditions and dsRNA-degrading nucleases in the saliva, hemolymph and gut has been associated to weak (or null) gene silencing response in other hemipterans, as *Acyrthosiphon pisum*^83^, *Lygus lineolaris*^84^ and *Myzus persicae*^85^.

Regarding *Dmai-BicC*, we were unable to identify any structures in the few eggs laid by *dsRNA^BicC^-*injected females, except for the presence of endosymbiont organisms, previously described in *D. maidis*^86–88^. Similar outcomes were achieved in the blood-sucking bug *Rhodnius prolixus*, suggesting that *BicC* role during hemipteran embryogenesis, could be prior to gastrulation^52^. Although most of the *dsRNA^BicC^-*fed females did lay eggs, we assign this phenomenon to the less amount of dsRNA consumed by these females, rather than a real difference in *Dmai-BicC* function. Unlike *D. melanogaster*, embryos presented no AP patterning defects^53,54^ and displayed the same phenotype as the control ones. *dsRNA^BicC^* microinjection into *D. maidis* females also resulted in a high percentage of sterility, agreeing with the disruption of *BicC* orthologs in *N. lugens*^51^ and *D. melanogaster*^55^ females.

Remarkably, the ovaries of *dsRNA^BicC^-*injected females shared some characteristics of atresia described in the bugs *R. prolixus*^89–91^ and *Dipetalogaster maxima*^92^, such as color changes in the cytoplasm of the oocytes and loss of columnar shape in some follicular cells. Follicular atresia is the mechanism by which the oocyte contents are degraded in response to stress conditions, allowing the energetic resources of the developing oocytes to be reallocated^93^. As this process has the ultimate goal of restoring female fitness and the most damaged ovaries corresponded to *dsRNA^BicC^-*fed females, it could be possible that the absence of some essential nutrients in the artificial diet may have contributed to an atretic-like phenotype, worsening the effects of *Dmai-BicC* silencing. In this sense, previous studies found that fertility was compromised in artificially-reared insects compared to those fed on plants^94,95^.

In summary, development of RNAi delivery methods, added to the availability of omics data will provide insights into the signaling networks controlling insect life cycle and propagation. In this regard, it is certain that *Dmai-BicC* is a key player in the reproduction of *D. maidis*.

## 5. CONCLUSION

This study provides evidence of a functional RNAi machinery in *D. maidis* and constitutes, to the best of our knowledge, the first report of RNAi in adult insects of this species. On the basis of our results, we propose that oogenesis-related genes, such as *Dmai-BicC*, are promising targets by inducing reproductive arrest of the females, hence preventing pest propagation. Considering both delivery methods, we believe that microinjection represents a robust technique for elucidating gene function, while oral administration could be the way forward in developing effective and sustainable RNAi-based control strategies against this leafhopper.

## 6. ACKNOWLEDGEMENTS

The authors thank the Centro de Investigaciones Básicas y Aplicadas (CIBA) for kindly allowing the use of their microscope facility.

