## Supplementary File for "Development of efficient RNAi methods in the corn leafhopper *Dalbulus maidis*, a promising application for pest control"

**Table S1. Primers used for dsRNA synthesis and RT-(q)PCR**

| Primer name | Usage | Sequence (5′–3′)+ | Amplicon length (bp) |
|---|---|---|---|
| *BicC_Fw* | RT-PCR | TAGAGCCGGCAGATGAGTTT | 372 |
| *BicC_Rv* | | TCCACACCTTCCATCTCTCC | |
| *BicC_Fw_T7* | dsRNA synthesis | **CGACTCACTATAGGG**TAGAGCCGGCAGATGAGTTT | 402 |
| *BicC_Rv_T7* | | **CGACTCACTATAGGG**TCCACACCTTCCATCTCTCC | |
| *BicC_Fw_qPCR* | RT-qPCR | ACCTGATGCTTCTTCACCCTAC | 128 |
| *BicC_Rv_qPCR* | | TAGCCAACTCCCATTCACATCC | |
| *Act1_Fw_qPCR* | RT-qPCR | CCGACAGGATGCAGAAGGAG | 219 |
| *Act1_Rv_qPCR* | | AGGGTTTGCGAGCTGAAGTT | |

+partial T7 promoter sequence is indicated in bold. T7 promoter: 5′-TAATA**CGACTCACTATAGGG**-3′.

### Figure S1

## (A)

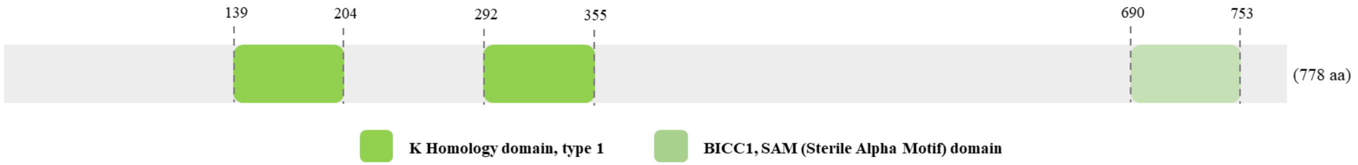


## (B)

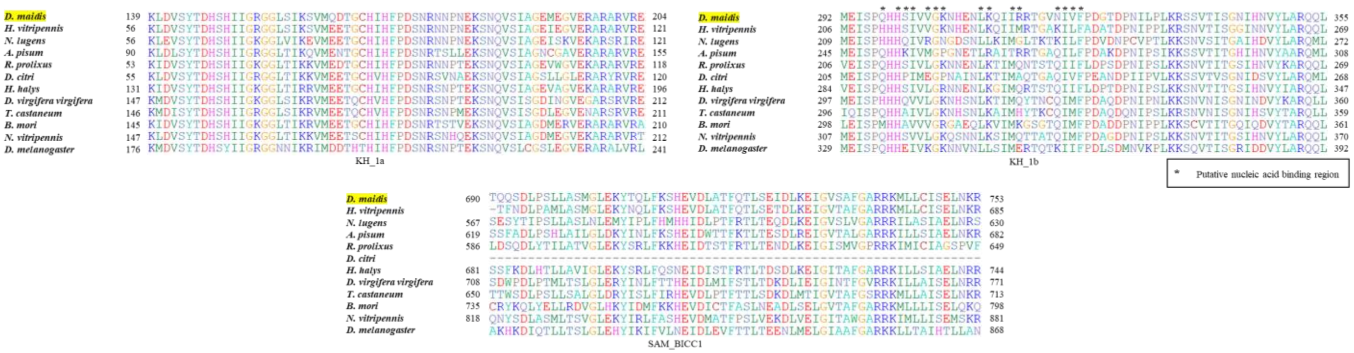


***In silico* characterization of *Dmai-BicC* protein**. **(A)** Diagram of *Dmai-BicC* predicted protein, including its conserved domains and length (in brackets). We identified TRINITY_DN24799_c0_g1_i7 as the putative *Dmai-BicC* transcript. Full length cDNA sequence was 2.655 bp long and encoded a 778 aa protein. **(B)** Partial alignment of *Dmai-BicC* predicted domains (KH_1a, KH_1b and SAM_BICC1) with orthologues from other species.

#### Figure S2

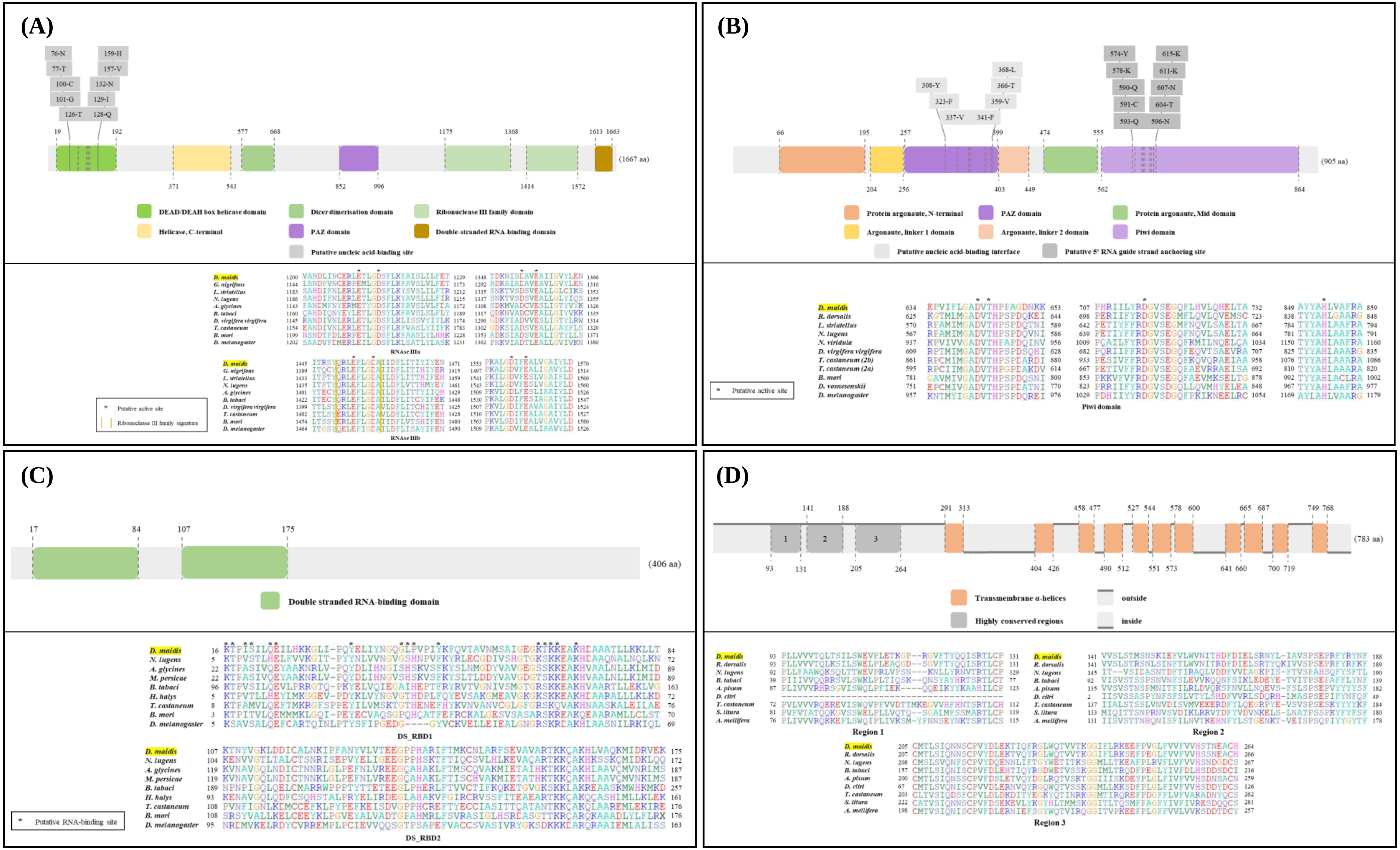


***In silico* characterization of RNAi-related genes in *D. maidis*.** Each panel shows a scheme of predicted *Dmai-Dcr2* **(A)** *Dmai-Ago2* **(B)**, *Dmai-R2D2* **(C)** and *Dmai-Sid1* **(D)** structure, including its domains and length in brackets (upper panel) and the partial alignments of one or more domain/s with orthologues from other species (lower panel). Asterisks show highly conserved residues.

#### Figure S3

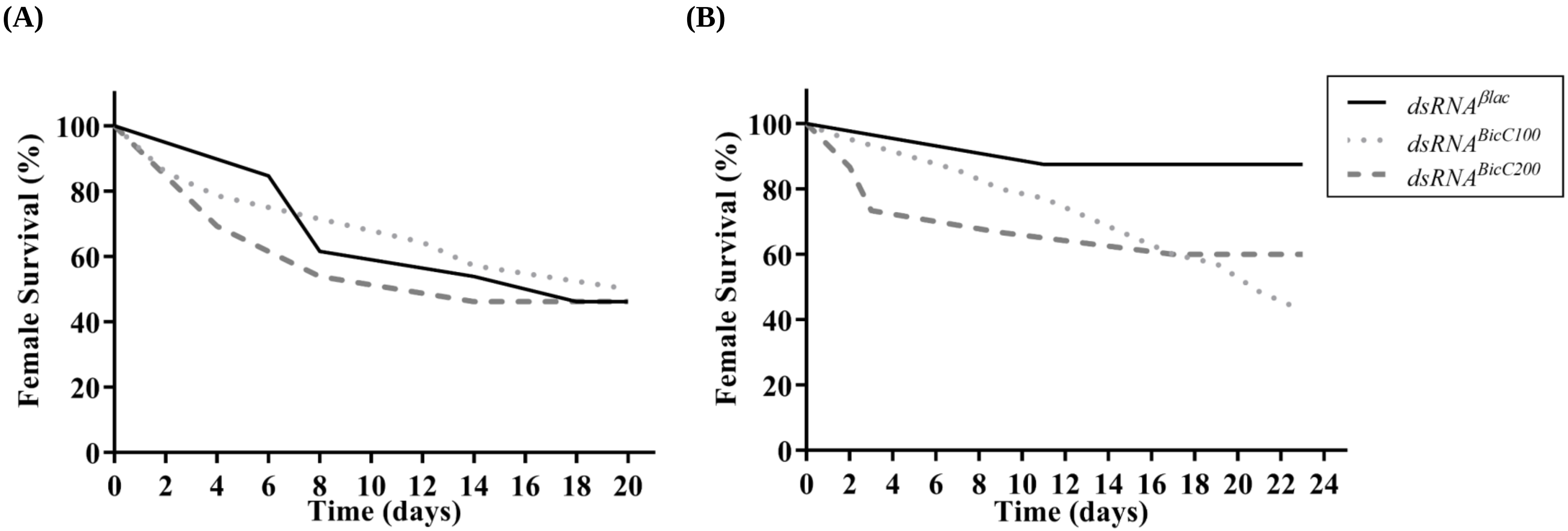


**Survival of *D. maidis* females after dsRNA delivery.** Percent survival of *D. maidis* females over the course of the injection- **(A)** and feeding-based **(B)** experiments. **(A)** 1: dsRNA microinjection day; 2: male introduction to the rearing cages. **(B)** 1: artificial diet feeding day; 2-4: dsRNA feeding days; 5: female and male introduction to the rearing cages. The survival curves were not significantly different according to Logrank test ($p \leq 0.05$).

#### Figure S4

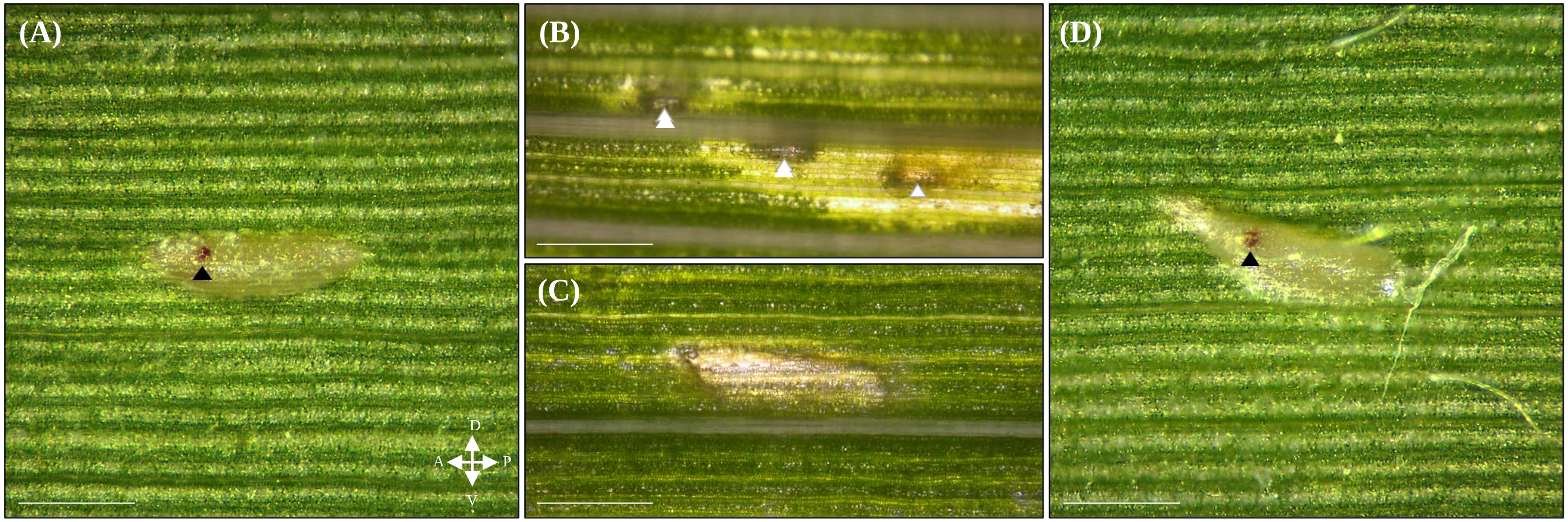


***In planta* ovipositions of *D. maidis* females after dsRNA delivery.** Eggs laid by control **(A)**, *dsRNA$^{BicC}$*-injected **(C)** and *dsRNA$^{BicC}$*-fed **(D)** females in maize leaves under dissecting microscope. Scale bar: 200 μm. Black arrowheads: red eye (very advanced embryo). White arrowheads: marks generated by the ovipositor apparatus as a sign of unsuccessful egg laying. A: anterior; P: posterior.

#### Figure S5

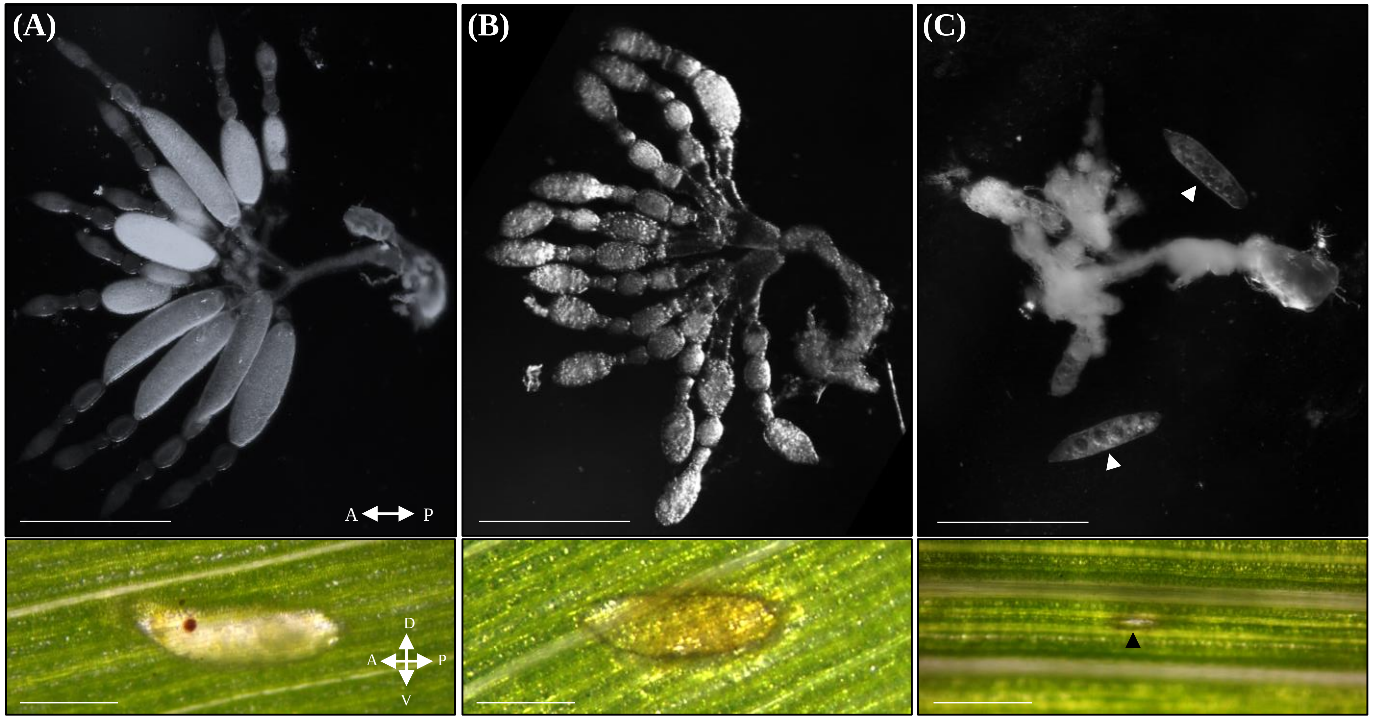


**Effect of *dsRNA$^{BicC}$* feeding on *D. maidis* females.** Ovary morphology of *dsRNA$^{BicC}$*-fed females (upper panel, scale bar: 500 µm) and their corresponding ovipositions (lower panel, scale bar: 200 µm) under dissecting microscope. Figures are ordered according to the severity of the phenotype, from **(A)** to **(C)**. White arrowheads: retained eggs dissected from the ovary for image capture. Black arrowheads: marks generated by the ovipositor apparatus as a sign of unsuccessful egg laying.
